# A K-mer based Bulked Segregant Analysis approach to map seed traits in unphased heterozygous potato genomes

**DOI:** 10.1101/2023.10.09.561609

**Authors:** Pajaree Sonsungsan, Mwaura Livingstone Nganga, Meric Lieberman, Kirk Amundson, Victoria Stewart, Kitiporn Plaimas, Luca Comai, Isabelle Henry

## Abstract

High-throughput sequencing-based methods for bulked segregant analysis (BSA) allow for the rapid identification of genetic markers associated with traits of interest. BSA studies have successfully identified qualitative (binary) and quantitative trait loci using QTL mapping. However, most traditional QTL mapping approaches require a reference genome. Here, we combine high throughput short read sequencing with bulk segregant analysis of k-mers (BSA-k-mer). This method can be applied to any organism and is particularly useful for species with genomes too different from the closest sequenced genome. It is also instrumental when dealing with highly heterozygous and polyploid genomes without phased haplotype assemblies and for which a single haplotype can control a trait. Finally, it is flexible in terms of population structure. Here, we present the application of the BSA-k-mer method for the rapid identification of candidate regions related to seed spot and seed size in diploid potato. While conventional QTL mapping of parental genotypes did not generate any signal, candidate loci were identified for each trait using the BSA-k-mer approach. The seed spot loci match with previously identified loci associated with pigmentation in potato. The loci associated with seed size are novel. Both sets of loci are potentially instrumental in future breeding towards true seeds in potato.

**Article Summary:** Identifying genes linked to agronomic traits in potatoes is challenging because potato genomes are complex and variable. We are investigating the genetic basis of seed size and color. Seeds were categorized as large or small, or spotted or not, based on simple visual observation. Next, DNA sequences from each individual were mined for association between random short sequences (k-mers) and those two traits. This more flexible method allowed us to identify regions of the potato genome associated with both traits.

## Introduction

Cultivated and natural populations of potato (*Solanum tuberosum*) display great phenotypic and genomic diversity (Hardigan *et al*. 2017), only part of which is captured by highly heterozygous commercial clones. Breeding is complicated by the autotetraploid genome of modern cultivated varieties. There is great interest in constructing inbred diploid parental varieties, and using these to produce F1 hybrids that can be distributed as botanical seed (Almekinders *et al*. 2009; Jansky *et al*. 2016). Central to this strategy is the use of genetic haploid inducers. These are diploids of the subspecies *phureja* that result in maternal (di)haploids when used to pollinate tetraploid cultivars (Hougas *et al*. 1958; Hermsen and Verdenius 1973). Haploid inducers share a common ancestry. They express a dominant anthocyanin marker, the embryo spot, which facilitates the early identification of dihaploids and true hybrids, by visual observation of the seed.

The presence of an embryo spot results from deposition of anthocyanins in the cotyledonary axils. Color in potatoes is a moderately complex trait determined by several loci that result in various color phenotypes (Zhang *et al*. 2009; Jung *et al*. 2009). Additionally, other loci govern color patterning. For example, color can be restricted to tuber eyes, tuber flesh, tuber skin, petals, floral abscission zone, nodes, stems, and abaxial or adaxial leaf surfaces (Ortiz and M Golmirzaie 2003; Zhang *et al*. 2009; Jung *et al*. 2009). It can also be expressed in the embryo, which results in a spot visible in the seed. Specifically, the presence of the embryo spot requires the action of two color genes: the presence of the B^d^ allele at regulator locus B, and either P or R. P and R result in purple and red colored anthocyanins, respectively, throughout the plant body (Dodds and Long 1955a; Endelman and Jansky 2016) and P is epistatic to R (Hermsen and Verdenius 1973). The presence of B^d^ is required for the deposition of anthocyanins specifically in the floral abscission zone, embryo spot, tuber eye spots and stem nodes (Dodds and Long 1955a). Haploid inducer IvP35 is homozygous for the embryo-spot genes (Hermsen and Verdenius 1973).

Another polymorphism of interest is seed size. Reliance on botanical seed requires that crosses between inbred parents produce seed that germinates and emerges readily after sowing. Seed size is also directly related to emergence, establishment and canopy size (Moles and Westoby 2004). The genetics of seed size has been investigated in the tomato, leading to the identification of several qtls (<u>Khan et al. 2012</u>), including the *Seed Weight 4.1* (or *Sw4.1*) locus located on chromosome 4, possibly associated with the function of an ABC transporter gene (<u>Orsi and Tanksley 2009</u>). In potato, many Quantitative Trait Loci (QTL) mapping and GWAS studies have been performed in both tetraploid and diploid populations to identify loci connected to many traits (Young 1996; Bradshaw *et al*. 2008; Byrne *et al*. 2020; Yuan *et al*. 2020; Prodhomme *et al*. 2020; Naeem *et al*. 2021; Alvarez-Morezuelas *et al*. 2023). However, there are no reports identifying genes or loci controlling seed size in potato.

To study the inheritance of both of these traits, we have hybridized diploid IVP35 (a haploid inducer strain) to a diploid, non-haploid inducer accession of *phureja* with contrasting traits. In the derived F1 and F2 populations we phenotyped color and seed size. To map these traits, we subjected 88 progeny to whole genome sequencing. Both parents displayed considerable heterozygosity complicating the analysis of haplotypes required for efficient mapping. To overcome this problem, we compared k-mers, arbitrary DNA sequences of k length, between bulked sets of progeny with different traits <u>(Michelmore et al. 1991; Sims et al. 2009;</u> <u>Nordström et al. 2013; Akagi et al. 2014; Prodhomme et al. 2020)</u>. In this case, the entire population was divided into two bulks since the two traits were binary and there were no extreme trait values. The method highlighted genomic regions involved in color and seed size determination that we could not readily identify by conventional mapping and the bulk strategy allows this analysis to be performed on a small population of individuals from different generations.

## Methods

### Plant materials

Potato clones IvP35 (PI 584995), which is homozygous for the embryo spot (purple) marker genes, and GND (PI 258855), which does not carry the purple spot marker, were obtained from the USDA Plant Germplasm center as *in vitro* plantlets and as seed, respectively. Both IVP35 and GND are diploid (2n = 2x = 24). Seeds were germinated and propagated aseptically in half-strength MS media, and *in vitro* plantlets were transplanted in the greenhouse. GND was crossed as the female parent to IvP35. Next, 52 of the resulting diploid F1 plants were grown in the greenhouse. To reduce chances of biased marker inheritance caused by the presence of less than four self-incompatibility (SI) alleles, pollen from these F1 plants was harvested and pooled, and used to pollinate emasculated flowers of the same pool of F1 plants. At least one berry from the intercrossed F1 flowers was collected from each of the F1 plants and its seeds collected to obtain the F2 lines.

### Seed germination and Crosses

Seeds that are at least three months old were germinated by soaking them in 1500 ppm GA3 (gibberellic acid) for 24-48 hours at room temperature to break any residual dormancy (Lam and Erickson 1966; Lam 1968). The seeds were then rinsed in water, and washed in soapy bleach solution (50% bleach with either 0.5% Tween-20 or Triton X-100) for up to 10 minutes. The seeds were rinsed in water and plated on half-strength MS media (0.5x MS with vitamins, 0.5% sucrose & 0.7% phytoagar). Germination was performed under cool and dark conditions where the highest germination rates were obtained (Lam and Erickson 1966; Lam 1968). Pollen was extracted by harvesting anthers just before or right after light browning at the tip. Anthers were placed on a filter paper and left to dry for 24-48 hrs. Pollen was then extracted using a vibrating rod (VegieBee™) on the folded filter paper, collected in 1.5 ml tubes and stored at 4°C or used immediately. Only pollen less than two weeks old was used for pollination. For pollination, female flowers were emasculated by removing the anthers at the start of yellowing. Pollen was then placed on the stigmas of these flowers either immediately or up to 3 days after emasculation, depending on flower maturity. Fruits were then harvested at around 30 days, and further ripened on the lab bench for up to 30 more days. Seeds were extracted from ripe fruits, rinsed with tap water, washed in 25% bleach for 10 minutes, rinsed, and placed on a filter paper to dry. The bleach wash also bleached the seed coat and increased visibility of the seed embryo spot. Seeds were stored for at least three months on the lab bench before being sown.

### Seed phenotyping

Both F1 and F2 seeds segregated in terms of size, and for the presence of the purple seed spot. Spotted and non-spotted seeds were identified visually simply by recording the presence or absence of the dark embryo spot. Seeds were divided into large (L) and small (S) bulks visually as well. In summary, each seed was assigned to one of four categories based on this very simple visual assessment: large spotted (LSP), large non-spotted (LNS), small spotted (SSP) and small non-spotted (SNS) after a rapid visual assessment.

### Sequence analysis

IvP35 was previously sequenced to around 30X coverage (Amundson *et al*. 2021). Leaf tissue from each of the F1 and F2 lines and GND were used for separate DNA extractions, library preparation, and sequencing using Novogene Corporation’s internal protocols. Paired-ended 150 bp reads were obtained on Illumina NovaSeq 6000 platform for an average of 10X read coverage per sample. Custom python scripts were used to demultiplex the reads and remove the adapter sequences (https://github.com/Comai-Lab/all prep). Independent libraries were obtained for each F1 and F2 individual. The list of samples sequenced can be found in (Supplementary Table S1).

### Trait mapping

Reads from high-throughput sequencing were mapped to the potato reference genome assembly version DM v6.1 (Pham *et al*. 2020) using the BWA mem algorithm (Li 2013) and default parameters. Variable positions were called using BCFtools version 1.10.2 (Danecek *et al*. 2021). Only reads with mapping quality greater than 30 were used and all duplicate variants removed (bcftools norm -d all). Aligned reads from GND and IvP 35 were compared to find genomic variants between the two parents. Sites with genotype inference score qualities lower than 100, or read depth higher than 70 (2.3x genomic average), or less than 25 (0.83x genomic average) were excluded. Positions that were homozygous for different alleles in IvP35 and GND were retained as parental SNPs (618,599 positions).

To genotype the F2 lines in terms of parental contributions, the genome was first partitioned into 100kb consecutive non-overlapping bins. For each individual, genotype was based on pooled information from all SNPs located with each bin. The minimum number of informative reads covering SNPs in a bin was set at 50. The genotype of bins for which fewer than 50 informative reads were available were left unassigned (NA). For bins with sufficient informative reads, a locus was called homozygous when more than 95% of reads were assigned to one parent, and heterozygous when more than 5% but less than 95% of the informative reads were assigned to both parental genotypes. These bins were used as markers for mapping using the qtl (r/qtl) package version 1.41-6 (Broman *et al*. 2003; Arends *et al*. 2010). Standard data cleanup outlined in the r/qtl manual was followed (Broman and Sen). Three mapping algorithms implemented in r/qtl were used including the EM algorithm, Haley-Knott regression, and multiple imputation (Dempster *et al*. 1977; Lander and Botstein 1989; Haley and Knott 1992; Chen 2005). The LOD threshold for significance was set by running permutation tests as recommended in the r/qtl manual.

### K-mer counting

Jellyfish, a multi-threaded hash based tool, was used to generate 31-bp k-mers present in our sequencing reads (Marçais and Kingsford 2011). This was performed independently for all samples. Next, k-mers counts were generated for each sample. k-mers that appeared only once in a sample were removed as they are likely to originate from sequencing errors or contamination. Next, the samples were divided into four phenotypic bulks (spotted seeds, non-spotted seeds, large seeds and small seeds), with each sample present in one of the seed size bulks and one of the seed color bulks. For each bulk, the k-mer lists from all individual samples were combined into a single list, containing k-mer sequences and counts for that bulk. Bulk k-mer profiles were generated using GenomeScope version 2.0. These profiles were used to identify a minimum k-mer count threshold for each bulk (Ranallo-Benavidez *et al*. 2020). Specifically, k-mers found less than 5 times in a given bulk were removed from the list. Finally, k-mer counts per bulk were merged into a single count table.

### Identification of significantly enriched k-mers

To identify k-mer that exhibited significantly different counts between two bulks, we applied a log likelihood ratio test for nested models (Rahman *et al*. 2018). Briefly, if we assume that each k-mer appears K_A_ times in bulk A and K_B_ times in bulk B, and that N_A_ and N_B_ are the total number of k-mers in bulk A and bulk B respectively, the k-mer counts are assumed to be Poisson distributed with rate K_A_/N_A_ and rate K_B_/N_B_ in bulk A and bulk B, respectively. The null hypothesis assumed that the rate of occurrence of a k-mer in bulk A and bulk B is not different. We first added 1 to all k-mer counts to avoid zero values. We used p-values to evaluate the significance of the chi-square statistics and performed a Bonferroni correction to account for multiple testing (Rahman *et al*. 2018). K-mers associated with adjusted p-values < 0.01 were retained as significantly enriched in one bulk. Finally, only k-mers exhibiting absolute log2 fold change > 2.5 between the two bulks were retained as differential k-mers. This resulted in a list of differentially enriched k-mers for each of the four bulks (Table 1).

**Table 1.**
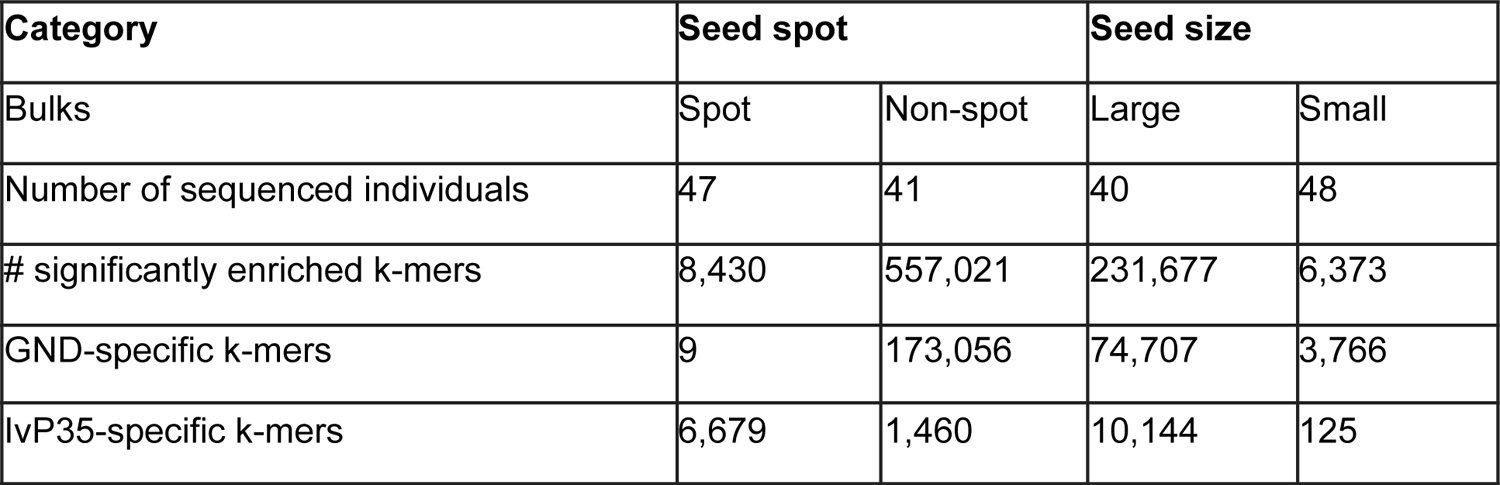
Number of significant k-mers in each bulk and how many were specific to each parent.

### Mapping and parent-of-origin of the enriched k-mers

All sequencing reads that contained at least one of the significant k-mers were identified. To visualize the location of the k-mers significant to each bulk, and their relative abundance, we performed dosage analysis as follows. Reads containing the significantly-enriched k-mers were mapped to the reference genome of the doubled monoploid potato of S. *tuberosum* Group Phureja DM v6.1 (Pham *et al*. 2020) using BWA version 0.7.17 (Li 2013). Next, we counted the number of reads mapping to each consecutive, non-overlapping 200 kb bin along the genome, for each bulk. Reads with mapping quality < 40 were discarded. This was performed using the custom python script bin-by-sam.py (https://github.com/Comai-Lab/bin-by-sam). To reduce bias due to difference in sample size in each bulk, we normalized the number of reads from the significant k-mers for each bulk to the total number of reads found in the corresponding bulk for each bin. To determine the parental origin of each k-mer, we compared k-mer abundance in the two parental readsets. A significantly enriched k-mers was classified as parent-specific if it was present in one parent but completely absent in the other. Reads containing these k-mers were mapped to the reference for localization, as described above.

### Data Availability

Sequence data have been deposited in the National Center for Biotechnology Information Sequence Read Archive: BioProject identifier PRJNA984282. Reads from parental accession IvP35 were previously deposited (<u>SRX10043416</u>).

## Results

In this study, two potato lines, ‘GND’ and ‘IvP35’, were crossed, and their F1 and F2 progeny analyzed to investigate the genetic factors associated with variation in seed size and seed spot. Both parental clones are highly heterozygous, making conventional mapping of all four haplotypes more challenging without the availability of phasing information between the parental haplotypes. Here, we combine a k-mer approach with bulk segregant analysis for the identification of bulk-specific sequences in potato.

### The F2 population varies in seed size and color

Clone GND (female), which does not exhibit a seed spot, was pollinated with IvP35 (male) which carries a seed purple spot, to generate progeny and observe seed traits. IvP35 exhibits deep purple flowers, flower/fruit abscission zone, tuber flesh and skin, internodes, and stem. GND, on the other hand, produces light purple flowers, white tuber flesh, white tuber skin, purple-red abaxial leaf surfaces, and lightly colored green stems.

Surprisingly, only approximately 57% of the F1 seed were spotted while the rest did not produce a seed spot. This is unexpected because IvP35 was bred by Hermsen & Verdenius to be homozygous for the dominant embryo spot factor (Hermsen and Verdenius 1973) (Singsit and Hanneman 1987). This suggests that genetic factors in GND might influence the expression of the embryo-spot trait.

Flowers from 52 F1 plants were intercrossed to produce seed, which we refer to as F2 from now on, for simplicity. The F2 seed fell into two size categories based on visual assessment (Figure 1). The F2 seed also fell into two classes based on the presence or absence of the embryo spot. There was no difference in seed size between spotted and non-spotted seeds. Of 1128 F2 seeds observed, 38.4% were spotted and 31% were small. All four phenotypic categories, large spotted (LSP), small spotted (SSP), large non-spotted (LNS), and small non-spotted (SNS) were represented. To investigate the genetic factors controlling seed size and seed color in this population, 88 samples were selected representing both large (N = 40) and small (N = 48), as well as spotted (N = 47) and non-spotted (N=41) seed (Table 1). These seeds were planted and tissue was collected from the germinated seedlings for DNA extraction and sequencing.

**Figure 1.**
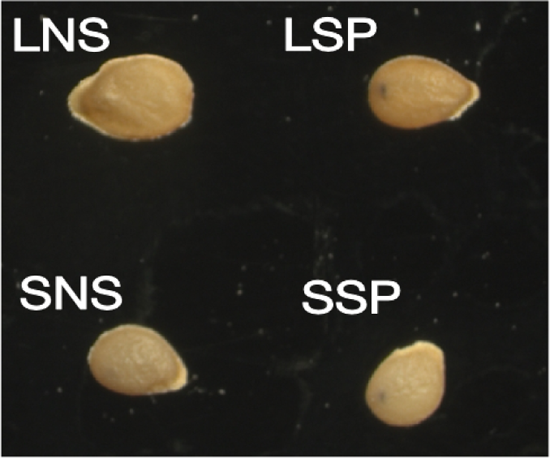
Seed traits. a) F1 and F2 Seeds were visually separated into four categories as follows: small spotted (SSP, bottom right), small non-spotted (SNS, bottom left), large non-spotted (LNS, top left), and large spotted (LSP, top right).

### Parental genotypes are not associated with either trait

To identify QTLs for the traits of interest, Illumina sequencing reads from a total of 88 F1 and F2 plants were aligned to the potato reference genome assembly version DM v6.1 (Pham *et al*. 2020). A total of 618,599 parental SNPs, which are homozygous in both parents but different between GND and IVP35, were used to characterize the F2 plants in terms of parental genotypes. SNP data were merged into 100 kb bins spanning the entire genome (see Methods for details). The percentage of heterozygous bins for the F2 individuals ranged between 32 and 78%, with a mean of 54%, as is expected from an F2 population, and confirming that no haploids were included in this F2 population.

We applied three methods to detect QTLs, including the EM algorithm, Haley-Knott regression, and multiple imputation (implemented in the r/qtl package). QTLs mapping using the parental SNPs did not identify any significant QTLs for either seed spot or seed size (results not shown). This is possibly due to the fact that a single parental haplotype controls these traits. For example, a segregating GND allele regulating the expression of the embryo seed spot would not easily be detected using homozygous parental genotypes. Haplotype-phased parental genotypes would be required to identify such factors, which is not available for this population.

### A k-mer-based identification of bulk-specific sequences without a reference sequence

To overcome this problem, we applied a k-mer-based method instead. For each trait, we split the population into two bulks and compared 31-mer k-mer count between pairs of bulks to identify k-mers associated with specific traits (Figure 2). The k-mer approach is computationally simple, may reduce biases in variant calling and, at least initially, does not require a reference genome. This method is thus more flexible and can in theory distinguish all four parental haplotypes.

**Figure 2.**
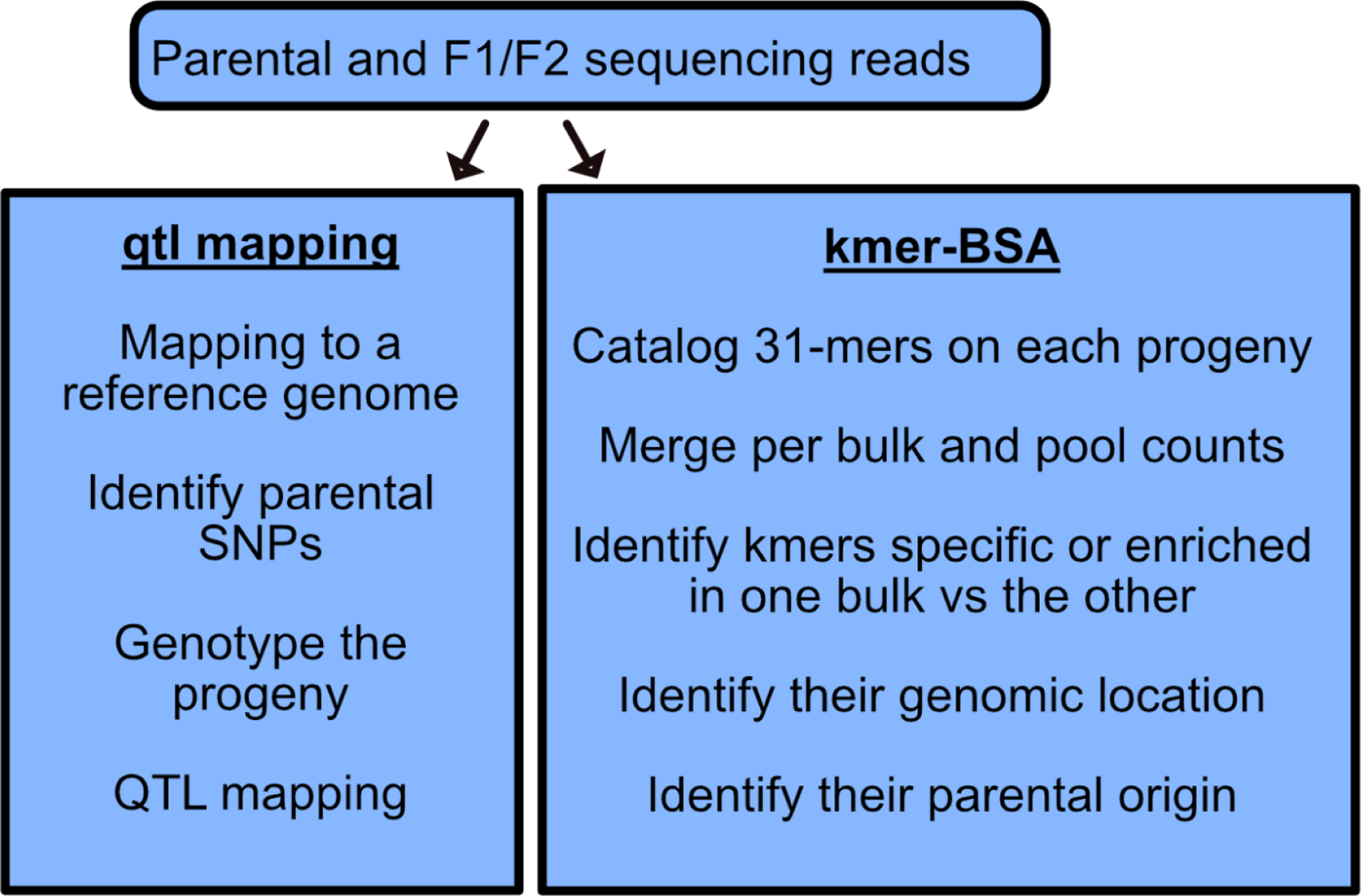
Overview of the two analysis pipelines. GND was pollinated with IvP35 to generate F1 plants, which were intercrossed to generate an F2 population. The plants were assigned to color and size categories based on visual observation of each seed. The F2 plants and a few F1 plants were sequenced individually using Illumina short-read sequencing. Trait mapping was performed using two approaches: QTL mapping based on each plant’s parental genotype, and BSA of k-mer counts. For the QTL analysis (left), Illumina short reads were mapped to the potato reference genome assembly and each individual was genotyped based on parental polymorphisms before QTL mapping. For the BSA-k-mer approach, Illumina short reads were used to generate 31-mer and k-mers were cataloged based on abundance in the two bulks for each trait. Significantly enriched k-mers were identified and traced back to the original sequencing reads, which were mapped to the reference sequence to identify regions of interest.

Figure 3 illustrates the principles underlying the identification of bulk-specific k-mers and highlights expectations, depending on the mode of action of loci responsible for these traits. For example, if we assume that large seed size is controlled by a single haplotype in one of the two parents (allele A), we can envision two simple situations. If the action of this allele is dominant, any F1 or F2 seed with at least one copy of A will be large while all others will be small. In terms of k-mers, we would expect that k-mers associated with the A alleles will be specific to the large bulk, and be readily identified by our pipeline. On the other hand, k-mers associated with the other three alleles (B, C and D), will be more abundant in the small bulk but present in both. Specifically, we expect those k-mers to be approximately 2.4x more abundant in the small bulk (14% vs 33%) and are less likely to be identified in our analysis since we are only retaining k-mers that exhibit a ≥ 2.5 fold enrichment in one bulk versus the other (see below). If the action of the A allele is recessive, only AA seed will be large. As a result, k-mers associated with the B, C and D alleles will be specific to the small bulk. k-mers associated with the A allele will be present in both bulks, but they will be significantly enriched in the large bulk compared to the small (100% vs 20% of alleles). They should therefore be identifiable in our analysis as well since we are retaining all k-mers with a 2.5 or higher enrichment in one bulk versus the other. In other words, loci associated with a dominant action are expected to be detected in only one of the two bulks while loci associated with a recessive allele are expected to be detectable in both bulks.

**Figure 3.**
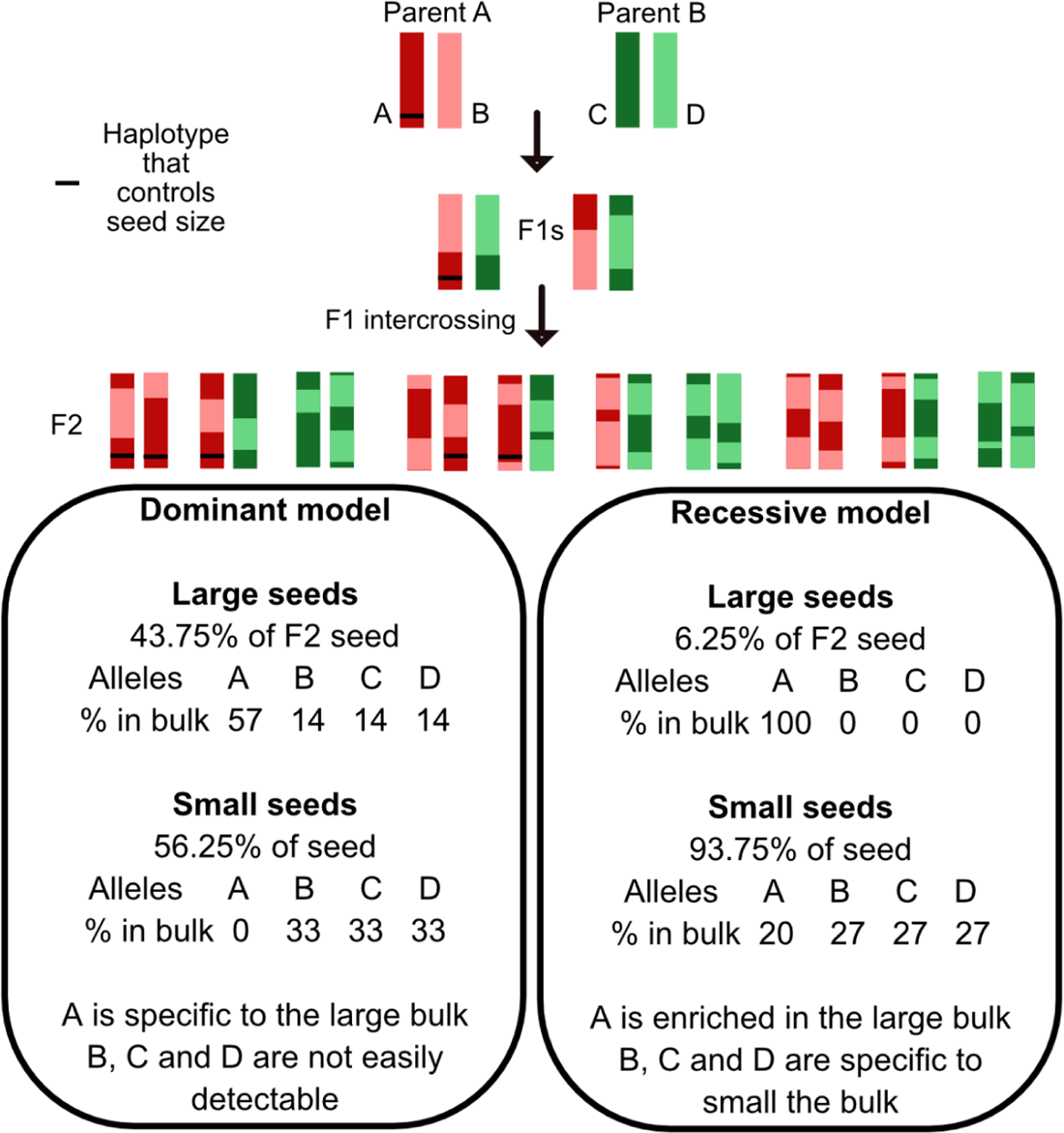
Expected k-mer enrichment patterns based on genetic mode of action of the causal loci. For any given locus controlling the trait, k-mer enrichment in one bulk or the other will vary depending on whether the allele acts recessively or in a dominant fashion. L represents the large seed genotype and S represents the small seed genotype.

Overall, between 2.5% and 4.15% of the initial k-mer sets were significantly enriched in one bulk versus the other (see Methods for details). Specifically, for the seed spot, we identified 565,451 k-mers that were significantly enriched in one bulk versus the other. Of those, 557,021 and 8,430 k-mers were enriched in the non-spotted bulk and the spotted bulk, respectively. For seed size, 238,050 significant k-mers were enriched in one bulk versus the other. Of those, 231,677 and 6,373 were significantly enriched in the large and small bulks, respectively. Based on the rationale presented above, these numbers suggest that the non-spotted and large seed traits could be controlled by a single dominant allele each.

### Mapping seed-color confirms previously identified QTLs

To characterize the location of the bulk-specific k-mers, we mapped the original reads containing these bulk-specific k-mers to a potato genomic assembly (version DM v6.1 (Pham *et al*. 2020)). We then derived the distribution of reads significantly enriched in the different bulks (Figures 4 and 5). Mapping the reads that contained k-mers significantly enriched in the spotted or non-spotted bulks resulted in the identification of two clear peaks in the non-spotted bulk, on chromosome 10 and 4 (Figure 4d). Specifically, one major peak was visible at the end of chromosome 10 (47.8 Mb to the end of the chromosome). Another high but very narrow peak was visible at the end of chromosome 4 (last 36 kb). The same two peaks were also visible in the location of the k-mers enriched in the spotted bulk but they were smaller (Figure 4a). Next, we determined whether these bulk-specific k-mers were specific to one parent or the other (Figure 4 b, c, e and f). When looking at the distribution of the parent-specific k-mers only, the same two peaks appeared again in the GND-specific k-mers but were absent in the IvP35-specific k-mers.

**Figure 4.**
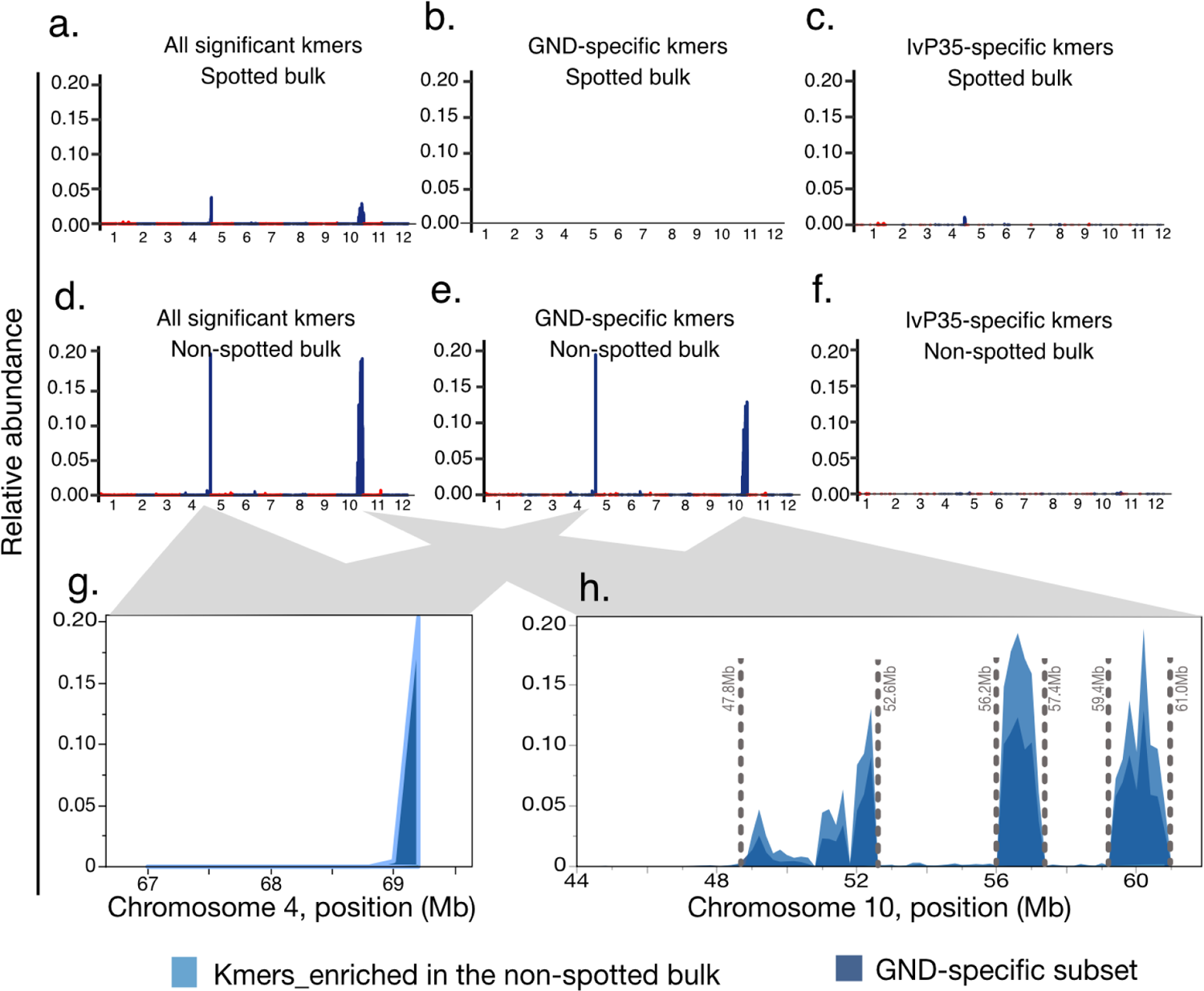
Mapping of the embryo spot loci. Distribution of k-mers significantly enriched in one bulk versus the other, depending on the bulk and parental origin. Taken together, the data presented here and the expectations described in Figure 3 suggest the existence of a GND-specific allele located on chromosome 10, which acts dominantly and sporophytically to prevent the expression of the purple spot in seed. (a,d) Mapping of reads containing enriched k-mers identified in the spotted or non-spotted seed bulks, respectively. For each plot, the number of reads per 250 kb bin relative to the total number of reads for the corresponding bulk is shown. (a, c, e) significant k-mers in the spotted seed bulk (b, d, f) significant k-mers in the non-spotted seed bulk. (a,b) all reads containing a significantly enriched k-mer (c-f) subset of the reads shown in (a and b) that are also parent-specific. (g,h) Detailed view of the peaks visible in (d) and (e). All k-mers enriched in the non-spotted bulk are in the background in light blue and those that are specific to the GND parent are overlaid on top in darker blue.

**Figure 5.**
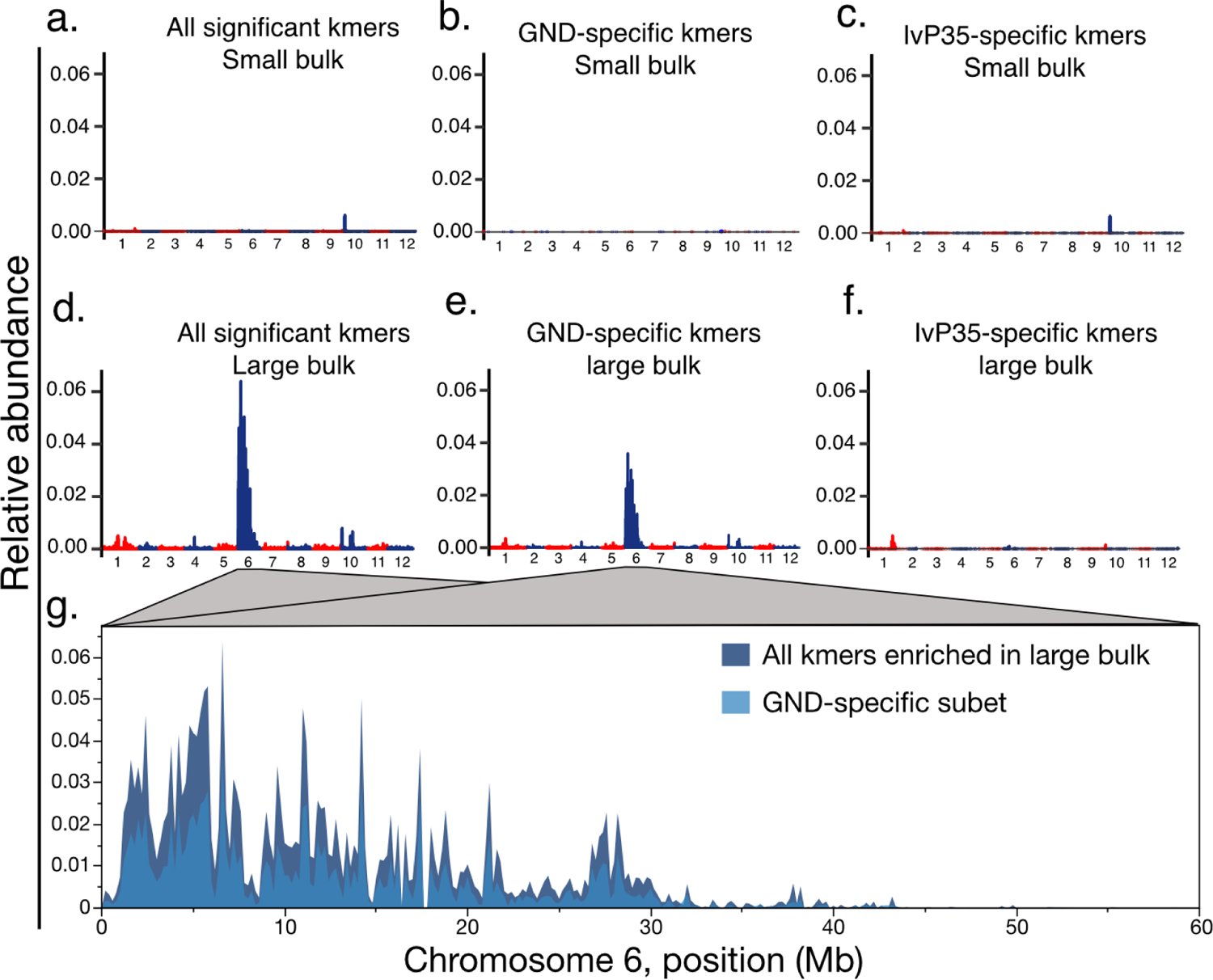
Mapping of seed size loci. Distribution of k-mers significantly-enriched in one bulk versus the other, depending on the bulk and parental origin. Taken together, the data presented here and the expectations described in Figure 3 suggest the existence of a GND-specific allele located on chromosome 6, which acts dominantly and sporophytically to produce large seed. (a,d) Mapping of reads containing enriched k-mers identified in the small or large seed bulks, respectively. For each plot, the number of reads per 250 kb bin relative to the total number of reads for the corresponding bulk is shown. (a, c, e) significant k-mers in the small seed bulk (b, d, f) significant k-mers in the large seed bulk. (a,b) all reads containing at least one significantly enriched k-mer (c-f) subset of the reads shown in (a and b) that are also parent-specific. (g,h) Detailed view of the peaks visible in (d) and (e). All k-mers enriched in the non-spotted bulk are in the background in light blue and those that are specific to the GND parent are overlaid on top in darker blue.

The peak on chromosome 4 is extremely narrow and located at the very distal end of the chromosome, suggesting that it is possibly associated with mis-assembly of the telomeric section of this chromosome, or mismapping of reads to the telomeric regions. For chromosome 10, the signal of significant k-mers spans a much wider region. Bulk-specific k-mer coverage varies widely across this region (48-60Mb). This dispersed signal may result from varying heterozygosity within GND, rather than recombination within this interval. Taken together, these results suggest that the allele underlying these peaks originated from GND. The identification of a color-associated locus on chromosome 10 is consistent with the previous identification of anthocyanin biosynthesis-related genes located on the distal arm of chromosome 10 (Endelman and Jansky 2016) and provides a clear proof-of-concept of the BSA-k-mer method in potato.

### Identification of genomic regions associated with seed size

We applied the same k-mer approach to identify loci associated with seed size. This resulted in the identification of a wide peak on chromosome 6, only present in the k-mers specific or enriched in the large bulk (Figure 5). Based on the percentage of large seeds observed in the F2 population (69% large seed), and the presence of a single peak in one bulk versus the other, our results are consistent with a dominant allele that confers increased seed size. Investigation of parental origin of these bulk-specific k-mers demonstrated that this dominant allele linked to large seed size originated from the GND parent (Figure 5).

## Discussion

Mapping traits of interest can be challenging if the reference genome is either not available or too divergent from the parental genomes at hand. Michelmore et al. (Michelmore *et al*. 1991) demonstrated that randomly-generated arbitrary PCR markers could be associated with polymorphic progeny bulks. The bulking approach became widely adopted with the advent of high-throughput sequencing technology. Using k-mers, Sims et al constructed bacterial phylogenies (Sims *et al*. 2009). Nordström et al identified mutations by bulking wild-type and mutant progeny (Nordström *et al*. 2013). Akagi identified and mapped the sex-determinant OGI comparing male and female cohorts of *Diospyros lotus* (Akagi *et al*. 2014). In the following years, multiple GWAS studies have employed k-mer analysis with success in systems with variable quality of reference genomes (Voichek and Weigel 2020; Kim *et al*. 2020). There are fewer examples of mapping natural variants in an experimental cross using k-mers, but the approach is extremely flexible and can be applied to many situations <u>(Prodhomme et al. 2020)</u>, such as the one presented here with individuals from both the F1 and F2 generation of a cross between two heterozygous parents Genome-agnostic association of k-mers to traits provides a reliable solution to the complexity of the pangenome (Vernikos *et al*. 2015): for example, a trait may depend on genes that are present in some population individuals, and absent in others and in the reference genome. In our case, we did not detect significant signals by using standard parent-based mapping approaches (see Methods), but instead were able to identify loci associated with both traits when detecting differential representations of k-mer counts between bulks. This speaks to the versatility of the k-mer-BSA approach. With no prior knowledge of the factor(s) regulating these traits or their potential mode of action, one cannot predict what type of cross will be most likely to enable their identification. The approach presented here has the potential to capture all model types and distinguish between all haplotypes.

Our analysis identified two regions associated with color, one on the distal arm of chromosome 10, and the other on the very distal end of chromosome 4. The peak on chromosome 4 is extremely narrow and appears fully telomeric. Specifically, there are no annotated genes past the 69.2 Mb location (where most of the bulk-specific k-mers are located) and only 5 genes in the less significantly enriched downstream bin (69.0 to 69.2 Mb). It is possible that this region of the genome is actually physically linked to the end of chromosome 10, but appears unlinked in the genomic reference that we are using, either because of an error in genome assembly, or because of a translocation events in GND compared to the reference from DM (*S. tuberosum* group Phureja).

The embryo purple spot in seed results from deposition of anthocyanins and its expression is controlled by a combination of genes that produce various color phenotypes (Dodds and Long 1955b; Jong 1991). The purple anthocyanins on embryo spots are determined by two dominant color genes, B and P (Dodds and Long 1955b; Endelman and Jansky 2016). Several previous mapping experiments have found the distal end of chromosome 10 to be associated with this phenotype. The region carries several anthocyanin pathway genes including two R2R3-MYB transcription factors and multiple anthocyanin biosynthesis genes (van Eck *et al*. 1994; Jung 2005; Jung *et al*. 2009; Endelman and Jansky 2016; Tengkun *et al*. 2019; Parra-Galindo *et al*. 2021).

The presence of the embryo seed spot is a critical component in the identification of haploids following crossing to haploid inducers from the Phurja group, such as IvP35. In typical tetraploid by diploid crosses, this method is usually reliable. But the results obtained here suggest that other natural alleles can suppress the expression of the IVP homozygous color alleles. Specifically, our data is consistent with the presence of a dominant and epistatic GND allele that regulates the action of the IVP embryo spot allele. As a result, we did obtain diploid individuals that carried the IvP35 allele, but did not exhibit the embryo spot. At first sight, there are two genes located in that region of chromosome 10 that can be identified as potential causal genes. The first gene is StAN1 (Soltu.DM.10G020850.1), a myb transcription factor, which controls anthocyanin levels (Bao *et al*. 2022). The second gene is StAN2 (Soltu.DM.10G020820.1), also a Myb-domain protein, previously shown to control the degree of anthocyanins in tuber (Jung *et al*. 2009; Zhang *et al*. 2020; Riveros-Loaiza *et al*. 2022). Identification of the causal allele and locus will help expand the use of haploid inducers to non-commercial potatoes.

Reads containing differential k-mers for seed size mapped to the proximal arm of chromosome 6, defining a very wide region of the chromosome. Many genes can contribute to seed size, acting either in maternal or zygotic tissues (Li *et al*. 2019). In our analysis, the differential reads had a distinct pattern: they originated from the GND parent and displayed enrichment predominantly in the large seed bulk. This could be explained if a GND allele enlarges seed in an additive or dominant manner. Further work fine-mapping this region will be necessary to identify candidate genes.

In summary, by detecting enrichment of k-mers derived from sequencing reads, we have mapped loci responsible for anthocyanin pigmentation and seed size. Discovery of a potential repressor of pigmentation GND allele on chromosome 10 provides a useful trait to breeders interested in developing true botanical seed in potato.

## Acknowledgements

This work was supported by the National Science Foundation Plant Genome Integrative Organismal Systems (IOS) Grant 2310230 to L.C. and I.M.H. PS was partially supported by the Research Assistantship Fund, Faculty of Science, Chulalongkorn University, and the Development and Promotion of Science and Technology Talents Project (Royal Government of Thailand scholarship).

